# Quantification of Enterohemorrhagic *Escherichia coli* O157:H7 proteome using TMT-Based Analysis

**DOI:** 10.1101/312652

**Authors:** Wanderson M. Silva, Jinlong Bei, Natalia Amigo, Pía Valacco, Ariel Amadio, Qi Zhang, Xiuju Wu, Ting yu, Mariano Larzabal, Zhuang Chen, Angel Cataldi

## Abstract

Enterohemorrhagic *Escherichia coli* (EHEC) O157:H7 is a human pathogen responsible for diarrhea, hemorrhagic colitis and hemolytic uremic syndrome (HUS). EHEC infection is distributed worldwide and numerous outbreaks of diseases caused by enterohemorrhagic have been reported. To promote a comprehensive insight into the molecular basis of EHEC O157:H7 physiology and pathogenesis, the combined proteome of EHEC O157:H7 strains, Clade 8 and Clade 6 isolated from cattle in Argentina, and the standard EDL933 (clade 3) strain has been analyzed. TMT (Tandem Mass Tags)-based quantitative proteomic and emPAI analyses were performed to estimate the protein abundance in EHEC proteome. 2,234 non-redundant proteins of EHEC O157:H7 were identified. A comparison of this result with *in silico* data of EHEC O157:H7 genome showed that approximately 40% of the predicted proteome of this pathogen were covered. According to the emPAI analysis, 85 proteins were among the most abundant (e.g. GAPDH, FliC H-antigen, Enolase, and GroEL). Tellurite resistance proteins were also highly abundant. COG analysis showed that although most of the identified proteins are related to cellular metabolism, the majority of the most abundant proteins are associated with translation processes. A KEGG enrichment analysis revealed that Glycolysis / Gluconeogenesis was the most significant pathway. On the other hand, the less abundant detected proteins are those related to DNA processes, cell respiration and prophage. Among the proteins that composed the Type III Secretion System, the most abundant protein was EspA. Altogether, the results show a subset of important proteins that contribute to physiology and pathogenicity of EHEC O157:H7.

**IMPORTANCE:** The study of the abundance of proteins present within a complex mixture of proteins in a cell, under different conditions, can provide important information about the activities of individual protein components and protein networks that are cornerstones for the comprehension of physiological adaptations in response to biological demands promoted by environmental changes. We generated a comprehensive and accurate quantitative list of EHEC O157:H7 proteome, which provides a description of the most abundant proteins produced by this pathogen that were related to physiology and pathogenesis of EHEC. This study provides information and extends the understanding on functional genomics and the biology of this pathogen.

## INTRODUCTION

Enterohemorrhagic *Escherichia coli* (EHEC) O157:H7 is a zoonotic pathogen belonging to Shiga toxin-producing *E. coli* (STEC) and responsible for different diseases as diarrhea, hemorrhagic colitis and hemolytic uremic syndrome (HUS). HUS is distributed worldwide and considered to be a public health problem in several countries (1,2). Unfortunately, Argentina is the country with the highest incidence of HUS in the world, with approximately 14 cases per 100,000 in children under 5 and a report of 500 cases per year (3,4). Cattle are the main reservoir of EHEC. Several studies have shown that most cases related to infection in human may be attributed to the high consumption of foods of bovine origin and especially ground beef is the main source of contamination (5).

Great efforts had been made to characterize strains of *E. coli* O157:H7 isolated from Argentinian cattle (6). Using the analysis of simple nucleotide polymorphisms, we have classified 16 strains of STEC O157:H7 in clade 6 and 8, which are the most virulent clades (6). *In vitro* and *in vivo* experimental results showed that the strains Rafaela II (clade 8) and 7.1 Anguil (clade 6) have a high virulence potential when compared with other strains and the standard strain EHEC O157:H7 EDL933 (7). These results enabled us to characterize the high prevalence of strains clade 6 and 8 in the Argentinian cattle. Importantly, these two clades might contribute to a high incidence of SUH in Argentina.

The availability of whole genome sequences of different EHEC strains has enabled genome-wide comparisons to identify factors that might be correlated to physiology and virulence of this pathogen (8). In addition, the implementation of system biology approaches, such as prediction of protein-protein network, has contributed substantially in the understanding of the pathogen and interactions with its host (9).

Information about the functions and activities of the individual proteins and pathways that control these systems is essential to understand complex processes occurring in living cells. Large scale quantitative proteomics is a powerful approach used to understand global proteomic dynamics in a cell, tissue or organism, and has been widely used to study protein profiles in the field of microbiology (10). Furthermore, the study of the abundance of proteins in different conditions or during different stages of growth or disease can provide important information about the activities of individual protein components or protein networks and pathways. The rapid growth of proteomic and genomic methods and tools has managed to reveal the basic protein inventory of a few hundred different organisms. Quantitative proteomic approaches have been applied to determine the absolute or relative abundance of proteins. This information gives insights about the biological function and properties of the cell as well as how cells respond to environmental or metabolic changes or stresses (11, 12). Quantitative proteomics analysis can contribute to the generation of datasets that are critical for our understanding of global proteins expression and modifications underlying the molecular mechanism of biological processes and disease states.

In a previous study, we reported the use of isobaric tags for comparative quantitation (TMT) method to identify the differentially expressed proteins among three EHEC O157:H7 isolates: Rafaela II (Clade 8), Anguil 7.1 (Clade 6) and EDL933 (Clade 3) (7). The proteome differences observed among these strains are related mainly to proteins involved in both virulence and cellular metabolism; which might reflect the virulence potential of each strain (7). The aim of the present study was to promote a more comprehensive insight into the molecular basis of EHEC O157:H7 physiology and pathogenesis. For this purpose, we applied high-throughput proteomics by performing TMT-based quantitative proteomic analysis and Exponential Modified Protein Abundance Index (emPAI) method (13) to quantify the EHEC O157:H7 proteome.

## RESULTS AND DISCUSSION

### Global proteomic analysis and functional classification of *Escherichia coli* (EHEC) O157:H7 proteome

In our proteomic analysis, we detected 2,519 non-redundant EHEC O157:H7 proteins (**Supplemental material, Table S.1**). When comparing this result with *in silico* data of EHEC O157:H7 genome, approximately 40% of the predicted proteome of this pathogen was identified (Figure 1A). To determine the abundance of the identified proteins, emPAI approach (13) was used. Because the emPAI value has a linear correlation with protein concentration, it allows a more accurate estimation of protein abundance. Considering only proteins with at least two peptides per protein, we quantified 2,234 proteins (**Supplemental material, Table S.2**). A dynamic range of protein abundance was generated spanning three orders of magnitude (Figure 1C). Eighty-five proteins were identified as most abundant in EHEC O157:H7 proteome (Table 1.) Of the 2,234 total proteins, 25 proteins are encoded by genes that are present in the pO157 plasmid; however, these proteins did not show a high abundance level **(Supplemental material, Table S.2)**.

**Figure 1:**
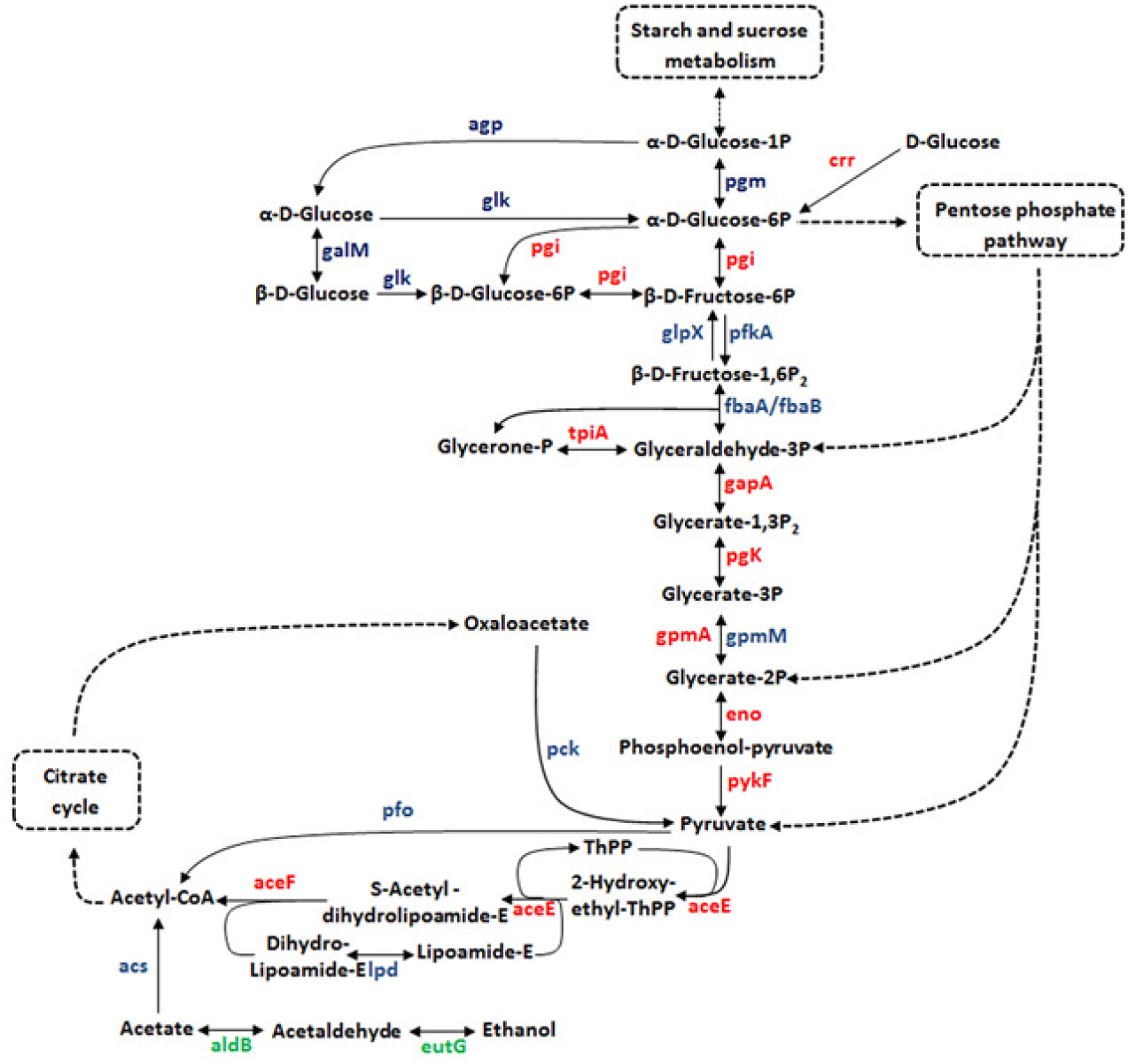
Characterization of EHEC O157:H7 proteome and correlation with *in silico* data. **(A)** Distribution of the peptides detected by MS. **(B)** Correlation of the proteomic results with *in silico* data of EHEC O157:H7 genome. **(C)** Dynamic range based on the emPAI value of the proteins identified by LC-MS analysis; pink, most abundant proteins; green, less abundant proteins and red, proteins that are present in the LEE pathogenicity island.

**Table 1:** List of the most abundant proteins of EHEC O157:H7 proteome

| Access Number | Description | Protein content (mol %) | COGSym bol<sup>(a)</sup> |
| --- | --- | --- | --- |
| gi\|667692306 | NAD-dependent glyceraldehyde-3-phosphate dehydrogenase | 3,71 | G |
| gi\|667694059 | SSU ribosomal protein S5p (S2e) | 3,55 | J |
| gi\|667692489 | Flagellar biosynthesis protein FliC | 3,38 | N |
| gi\|667693503 | Enolase | 3,27 | G |
| gi\|667693720 | putative Fe(2+)-trafficking protein YggX | 3,17 | PO |
| gi\|667693013 | Hypothetical protein | 3,09 | S |
| gi\|667695021 | Heat shock protein 60 family chaperone GroEL | 3,08 | O |
| gi\|667689612 | Chaperone protein DnaK | 3,04 | O |
| gi\|667690667 | Tellurium resistance protein TerD | 3,00 | T |
| gi\|667693130 | Phosphotransferase system, phosphocarrier protein | 2,85 | TG |
| gi\|667694476 | EspA | 2,82 | J |
| gi\|667694778 | Triosephosphate isomerase | 2,79 | G |
| gi\|667691933 | Thiol peroxidase, Tpx-type | 2,78 | O |
| gi\|667692763 | DNA-damage-inducible protein I | 2,78 | P |
| gi\|667694337 | Dipeptide-binding ABC transporter | 2,76 | E |
| gi\|667690668 | Tellurium resistance protein TerE | 2,75 | T |
| gi\|667694081 | Translation elongation factor Tu | 2,69 | J |
| gi\|667690420 | Phosphoglycerate mutase | 2,68 | G |
| gi\|667693674 | Phosphoglycerate kinase | 2,65 | G |
| gi\|667691384 | Isocitrate dehydrogenase [NADP] | 2,64 | C |
| gi\|667689773 | Translation elongation factor Ts | 2,63 | J |
| gi\|667695123 | Endoribonuclease L-PSP | 2,62 | V |
| gi\|667691519 | DNA-binding protein H-NS | 2,61 | K |
| gi\|667691364 | Hypothetical protein | 2,59 | I |
| gi\|667691726 | Glutamate decarboxylase | 2,56 | E |
| gi\|667690418 | putative secreted protein | 2,56 | S |
| gi\|667694844 | LSU ribosomal protein L1p (L10Ae) | 2,54 | J |
| gi\|667694305 | Glutamate decarboxylase | 2,52 | E |
| gi\|667695075 | SSU ribosomal protein S6p | 2,47 | J |
| gi\|667690270 | Alkyl hydroperoxidoreductase protein C | 2,46 | V |
| gi\|667695256 | Purine nucleoside phosphorylase | 2,45 | F |
| gi\|667689607 | Transaldolase | 2,43 | G |
| gi\|667695020 | Heat shock protein 60 family co-chaperone GroES | 2,43 | O |
| gi\|667694296 | Chaperone HdeA | 2,42 | C |
| gi\|667694705 | Periplasmic thiol:disulfide interchange protein | 2,41 | O |
| gi\|667690756 | Phosphoserine aminotransferase | 2,39 | HE |
| gi\|667694406 | Glutaredoxin 3 (Grx3) | 2,39 | O |
| gi\|667692485 | Cystine ABC transporter | 2,38 | ET |
| gi\|667692357 | Cold shock protein CspA | 2,37 | K |
| gi\|667694062 | SSU ribosomal protein S8p (S15Ae) | 2,34 | J |
| gi\|667694071 | LSU ribosomal protein L22p (L17e) | 2,33 | J |
| gi\|667695077 | LSU ribosomal protein L9p | 2,32 | J |
| gi\|667691029 | Ferrous iron transport periplasmic protein | 2,29 | C |
| gi\|667691774 | Hypothetical protein | 2,29 | R |
| gi\|667694592 | ATP synthase beta chain | 2,29 | C |
| gi\|667694268 | Universal stress protein A | 2,29 | T |
| gi\|667690733 | Translation initiation factor 1 | 2,27 | J |
| gi\|667694868 | IMP cyclohydrolase | 2,27 | F |
| gi\|667694766 | Manganese superoxide dismutase | 2,26 | P |
| gi\|667690203 | Peptidyl-prolyl cis-trans isomerase PpiB | 2,25 | O |
| gi\|667694064 | LSU ribosomal protein L5p (L11e) | 2,25 | J |
| gi\|667690810 | Outer membrane protein A precursor | 2,24 | M |
| gi\|667691522 | Alcohol dehydrogenase | 2,24 | C |
| gi\|667693128 | Cysteine synthase | 2,24 | E |
| gi\|667690777 | Aspartate aminotransferase | 2,24 | E |
| gi\|667694846 | LSU ribosomal protein L7/L12 (P1/P2) | 2,23 | J |
| gi\|667690752 | Pyruvate formate-lyase | 2,23 | C |
| gi\|667693266 | Serine hydroxymethyltransferase | 2,22 | E |
| gi\|667691738 | Osmotically inducible protein C | 2,21 | V |
| gi\|667689716 | Dihydrolipoamide acetyltransferase | 2,20 | C |
| gi\|667693332 | DNA-damage-inducible protein I | 2,19 | OT |
| gi\|667689715 | Pyruvate dehydrogenase E1 component | 2,16 | C |
| gi\|667690091 | hypothetical protein YajQ | 2,16 | S |
| gi\|667690760 | SSU ribosomal protein S1p | 2,15 | K |
| gi\|667694072 | SSU ribosomal protein S19p (S15e) | 2,14 | J |
| gi\|667692196 | Pyruvate kinase | 2,14 | G |
| gi\|667691439 | Protease VII (OmpT) precursor | 2,14 | M |
| gi\|667693243 | Iron-sulfur cluster assembly scaffold protein IscU | 2,13 | O |
| gi\|667692658 | 6-phosphogluconate dehydrogenase, decarboxylating | 2,12 | G |
| gi\|667693132 | PTS system, glucose-specific IIA component | 2,11 | G |
| gi\|667694683 | Uridine phosphorylase | 2,11 | F |
| gi\|667689775 | Ribosome recycling factor | 2,11 | J |
| gi\|667690429 | Molybdenum ABC transporter | 2,10 | P |
| gi\|667694065 | LSU ribosomal protein L24p (L26e) | 2,10 | J |
| gi\|667689933 | Hypothetical protein | 2,08 | F |
| gi\|667694594 | Peptidyl-prolylcis-trans isomerasePpiA precursor | 2,07 | C |
| gi\|667694106 | ATP synthase alpha chain | 2,07 | O |
| gi\|667690289 | Cold shock protein CspA | 2,06 | K |
| gi\|667695103 | Inorganic pyrophosphatase | 2,06 | CP |
| gi\|667694625 | Ketol-acid reductoisomerase | 2,06 | EH |
| gi\|667694890 | Glucose-6-phosphate isomerase | 2,05 | G |
| gi\|667690141 | hypothetical protein co-occurring with RecR | 2,03 | R |
| gi\|667694076 | LSU ribosomal protein L3p (L3e) | 2,03 | J |
| gi\|667693201 | Arsenate reductase | 2,03 | P |
| gi\|667694068 | LSU ribosomal protein L29p (L35e) | 2,01 | J |

(a) COG groups are defined in the legend to Fig. 2B.

The emPAI method was applied to estimate the relative quantification of the EHEC O157:H7 proteome from a dataset generated with strains *E. coli* O157:H7 Rafaela II, Anguil 7.1 and EDL933. The strains were grown in D-MEM media and then, proteins from total bacterial lysates were extracted and digested in solution. The resulting peptides were labeled with isobaric reagents. Finally samples were pooled and peptides were analyzed by 2D-LC MS/MS.

The emPAI method was applied to estimate the relative quantification of the EHEC O157:H7 proteome from a dataset generated with strains *E. coli* O157:H7 Rafaela II, Anguil 7.1 and EDL933. The strains were grown in D-MEM media and then, proteins from total bacterial lysates were extracted and digested in solution. The resulting peptides were labeled with isobaric reagents. Finally samples were pooled and peptides were analyzed by 2D-LC MS/MS.

We subsequently performed functional annotation of the identified proteins using gene ontology (14). Cluster of orthologous group (COG) analysis grouped the identified proteins into four important functional groups: (i) metabolism, (ii) information storage and processing, (iii) cellular processes and signaling, and (iv) poorly characterized (Figure 2A). Although most of the identified proteins are related to cellular metabolism, the most abundant proteins are involved in the translation process, followed by energy metabolism and posttranslational modification, protein turnover and chaperones, which shows an intense metabolic activity mainly in the protein synthesis (Figure 2B). On the other hand, most of the less abundant proteins are involved in replication, recombination and repair (Figure 2B).

**Figure 2:**
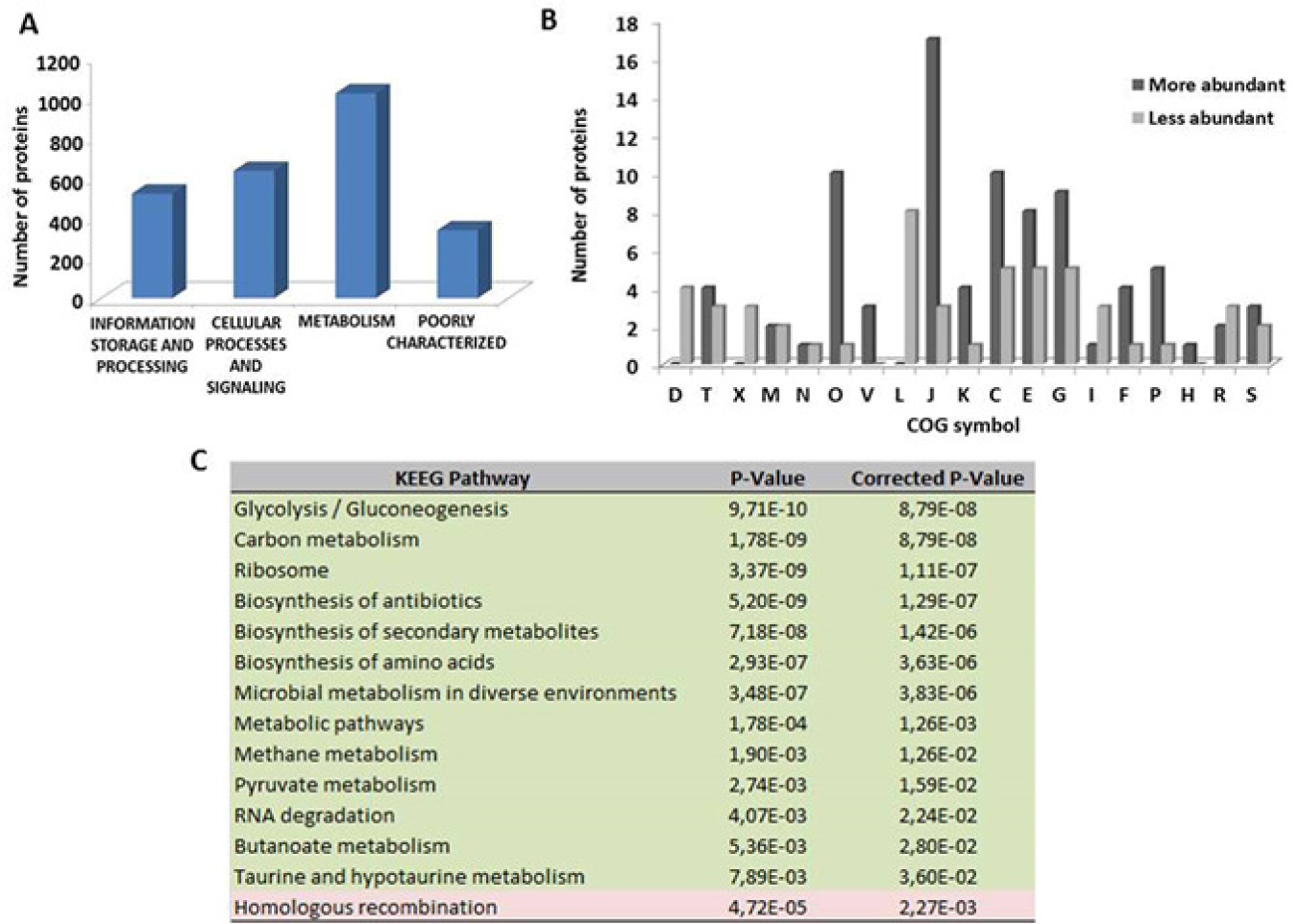
Functional analysis of the EHEC O157:H7 proteome. **(A)** Proteins classified by COG functional categories **(B)** Categorization of the proteins identified into biological processes. [E] Amino acid transport and metabolism; [G] carbohydrate transport and metabolism; [D] cell cycle control, cell division, and chromosome partitioning; [N] cell motility; [M] cell wall/membrane/envelope biogenesis; [H] coenzyme transport and metabolism; [V] defense mechanisms; [C] energy production and conversion; [W] extracellular structures; [S] function unknown; [R] general function prediction only; [P] inorganic ion transport and metabolism; [U] intracellular trafficking, secretion, and vesicular transport; [I] lipid transport and metabolism; [X] mobilome: prophages, transposons; [F] nucleotide transportand metabolism; [O] post-translational modification, protein turnover and chaperones; [L] replication, recombination, and repair; [A] RNA processing and modification; [Q] secondary metabolite biosynthesis, transport, and catabolism; [T] signal transduction mechanisms; [K] transcription; [J] translation, ribosomal structure and biogenesis. **(C)** KEGG pathway enrichment analysis, the colors are based on the protein abundance; blue, most abundant and green, less abundant.

Pieper et al. (15) and Ishihama et al. (16) also conducted proteomic studies on *E. coli* K-12 and EHEC O157:H7 strain 86-24, respectively, to determine the absolute abundance of proteins. Seventy proteins of the most abundant proteins in our study were also found as the most abundant proteins in *E. coli* K-12 **(Table 2)**. Those proteins are related to carbohydrate metabolism, transcription, translation, posttranslational modification and signal transduction mechanisms (16). On the other hand, only 12 proteins of the most abundant group **(Table 2)** were the most abundant ones in the data obtained from quantitative proteome of EHEC O157:H7 strain 86-24 (15). Some of those proteins (e.g. TerD, TerE, EspA and DNA-damage-inducible protein I) are absent from *E. coli* K-12. Interestingly, when comparing our results with those of Pieper et al. (15) and Ishihama et al. (16), the *E. coli* proteome was evaluated in different grown condition. Despite the different growth conditions, glyceraldehyde-3-phosphate dehydrogenase, translation elongation factor Tu, DNA-binding protein H-NS, alkyl hydroperoxidereductase protein C, GroEL chaperone and 50S ribosomal protein L7/L12 were detected as the most abundant proteins as well (Table 2). These results suggest a set of proteins that may play an important role in the biology of *E. coli*.

**Table 2:**
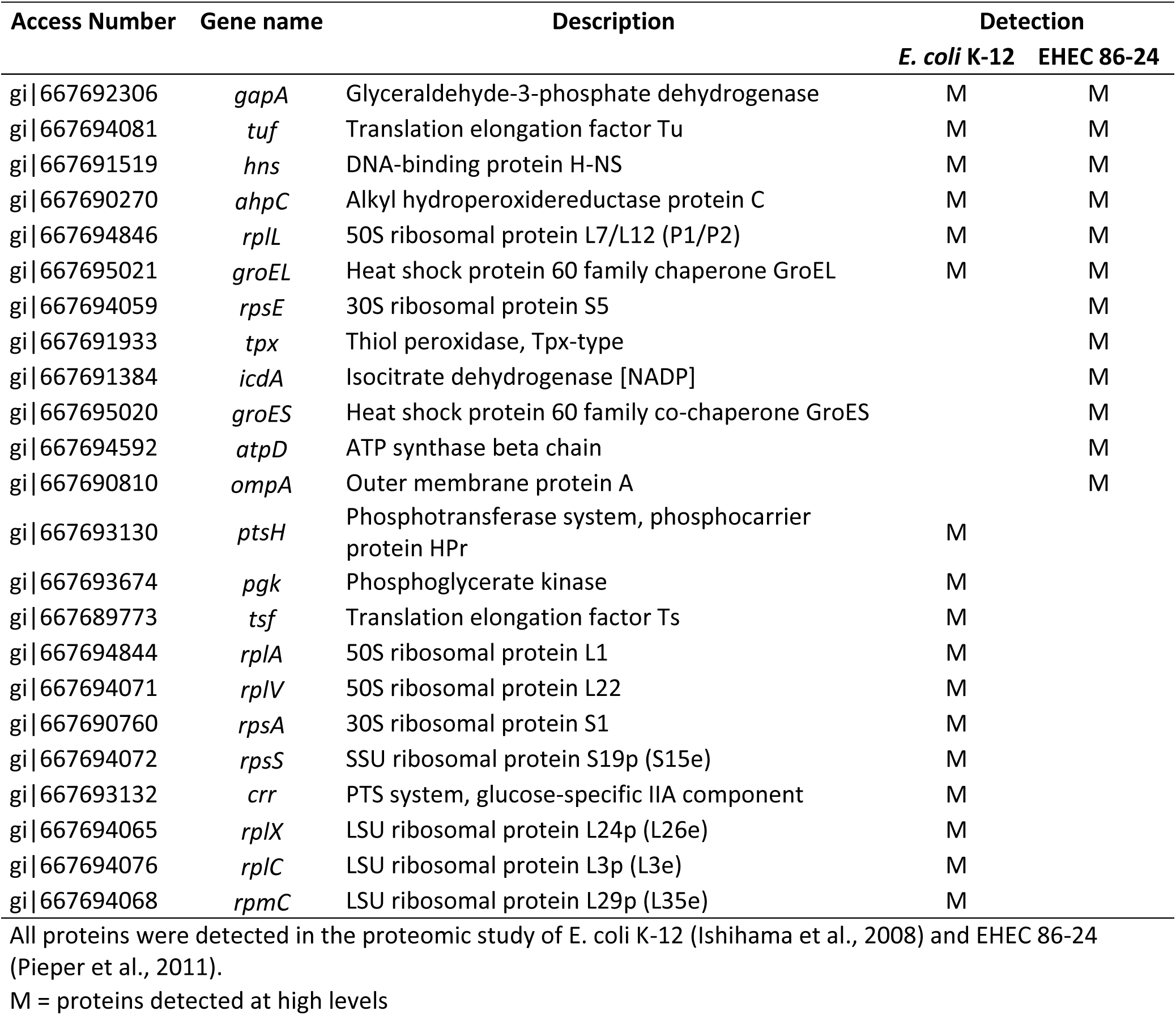
List of the most abundant proteins detected in *E. coli* K-12 and EHEC 86-24

| Access Number | Gene name | Description | Detection |  |
| --- | --- | --- | --- | --- |
|  |  |  | *E. coli* K-12 | EHEC 86-24 |
| gi\|667692306 | *gapA* | Glyceraldehyde-3-phosphate dehydrogenase | M | M |
| gi\|667694081 | *tuf* | Translation elongation factor Tu | M | M |
| gi\|667691519 | *hns* | DNA-binding protein H-NS | M | M |
| gi\|667690270 | *ahpC* | Alkyl hydroperoxidoreductase protein C | M | M |
| gi\|667694846 | *rplL* | 50S ribosomal protein L7/L12 (P1/P2) | M | M |
| gi\|667695021 | *groEL* | Heat shock protein 60 family chaperone GroEL | M | M |
| gi\|667694059 | *rpsE* | 30S ribosomal protein S5 |  | M |
| gi\|667691933 | *tpx* | Thiol peroxidase, Tpx-type |  | M |
| gi\|667691384 | *icdA* | Isocitrate dehydrogenase [NADP] |  | M |
| gi\|667695020 | *groES* | Heat shock protein 60 family co-chaperone GroES |  | M |
| gi\|667694592 | *atpD* | ATP synthase beta chain |  | M |
| gi\|667690810 | *ompA* | Outer membrane protein A |  | M |
| gi\|667693130 | *ptsH* | Phosphotransferase system, phosphocarrier protein HPr | M |  |
| gi\|667693674 | *pgk* | Phosphoglycerate kinase | M |  |
| gi\|667689773 | *tsf* | Translation elongation factor Ts | M |  |
| gi\|667694844 | *rplA* | 50S ribosomal protein L1 | M |  |
| gi\|667694071 | *rplV* | 50S ribosomal protein L22 | M |  |
| gi\|667690760 | *rpsA* | 30S ribosomal protein S1 | M |  |
| gi\|667694072 | *rpsS* | SSU ribosomal protein S19p (S15e) | M |  |
| gi\|667693132 | *crr* | PTS system, glucose-specific IIA component | M |  |
| gi\|667694065 | *rplX* | LSU ribosomal protein L24p (L26e) | M |  |
| gi\|667694076 | *rplC* | LSU ribosomal protein L3p (L3e) | M |  |
| gi\|667694068 | *rpmC* | LSU ribosomal protein L29p (L35e) | M |  |

All proteins were detected in the proteomic study of *E. coli* K-12 (Ishihama et al., 2008) and EHEC 86-24 (Pieper et al., 2011).
M = proteins detected at high levels

We also detected shiga-toxin subunits such as StxA, StxB, Stx2a and Stx2cb; these proteins, however, were not among the most abundant proteins (Figure 1). Pieper et al. (15) also obtained similar results in EHEC 86-24 proteome. This low abundance can be associated with environmental or nutritional conditions that contribute to the bacterial lysis and consequently to the production of the toxin (15, 17, 18).

### Metabolic Network Analysis

To identify the most relevant biological pathways of the identified proteins, we performed a KEGG enrichment analysis. This analysis provides a comprehensive understanding about pathways that might contribute to cellular physiology (19). When we evaluated the most abundant proteins, we identified 13 pathways that were considered significant (*p* < 0.05) and the Glycolysis / Gluconeogenesis was the most significant (Figure 2C). On the other hand, among the less abundant proteins, we only detected the homologous recombination pathway, which is extremely important for the accurate repair of DNA double-strand breaks (Figure 2C). Different studies have reported that glycolysis / gluconeogenesis pathway might influence in the colonization process of EHEC in the gastrointestinal tract of both mouse and bovine (20, 21). Although glycolysis substrates inhibit the expression of genes that are localized in locus of enterocyte effacement (LEE), this pathway plays an important role in the initial colonization and maintenance of EHEC in the mouse intestine. In addition, gluconeogenesis not only induces LEE gene expression, but contributes to the later stages of EHEC colonization in mouse as well (20, 22).

In our proteomic analysis, 23 proteins that composed the Glycolysis / Gluconeogenesis pathway of *E. coli* were identified (Figure 3). NAD-dependent glyceraldehyde-3-phosphate dehydrogenase (GAPDH) was the most abundant protein of EHEC O157:H7 proteome (Table 1). This important cytoplasmic protein of the Glycolysis pathway is also described as a moonlight protein, owing to the distinct functions performed by this enzyme in different cellular localization (23). Some studies showed that GAPDH secreted by EHEC and enteropathogenic *E. coli* (EPEC) strains can bind to fibrinogen and epithelial cell, which could contribute to the pathogenesis of this bacterium mainly through cell adhesion (24, 25). Another protein that is also described as a moonlight protein and was detected among the most abundant proteins of the EHEC proteome is enolase (Table 1) (23). This glycolytic enzyme that plays an important role in the carbon metabolism also acts in the RNA degradosome process, mainly in the RNA processing and gene regulation. In *E. coli*, enolase-RNase E/ degradosome complex regulates bacterial morphology under anaerobic condition by inducing a filamentous form, which is observed by some pathogenic *E. coli* strains under oxygen limiting conditions (26).

**Figure 3:**
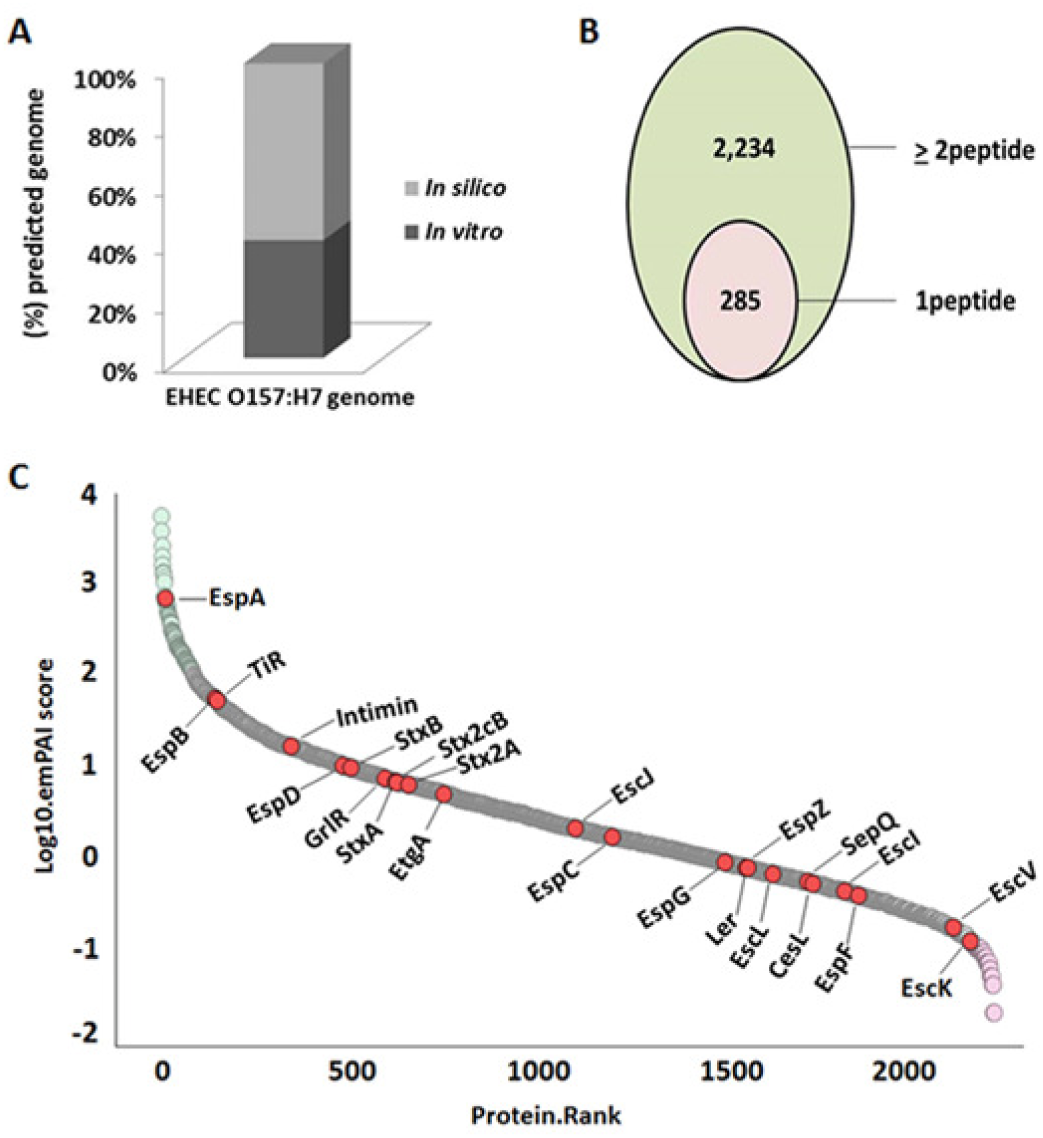
Overview of the Glycolysis / Gluconeogenesis pathway of EHECO 157:H7. Enzymes of the Glycolysis / Gluconeogenesis metabolism that were identified at the proteome level. Blue, proteins detected in our proteomic analysis; Green, proteins not identified in our study and Red, proteins detected as most abundant.

### Information storage and processing

Most proteins described as the most abundant are involved in translation processes. Similar results had been observed in *E. coli* K-12 (15). In addition, according to the KEGG enrichment analysis, the ribosome was strongly enriched (Figure 2C). We identified proteins involved in structural elements of the ribosome as well as related to initiation, elongation and terminations steps, which are required to the translation process (27). Among these proteins, the translation elongation factor Tu was identified (EF-Tu) (Table 1). EF-Tu could play a role in the resistance process of this bacterium in the gastrointestinal tract (28), as well as against cellular damage generated by bile salt sodium deoxycholate (29). These results show an intense metabolic activity of EHEC mainly in protein synthesis. Unlike *E. coli* K-12 (15), the proteins involved in transcription process in EHEC were identified as most abundant. CspA was identified to be among the most abundant proteins as well. This RNA chaperone is described as the major cold shock protein of *E. coli*. CspA binds to RNA molecules and destabilizes stem loop structures to prevent and resolve misfolding of RNA (30).

### Cellular processes and signaling

Flagella are filamentous structures that contribute to pathogenesis of pathogenic *E. coli*, mainly in motility, adhesion and biofilm production (31). Generally, this organelle is constituted by basal body, hook and a filament that is composed by flagelin or flagellar antigen FliC, which belongs to the H-antigens group (31, 32). FliC was detected as highly abundant (Table 1). In addition, a study performed in EPEC showed that FliC might be involved in the inflammatory response during the EPEC infection, due to the capacity of flagelin to induce interleukin-8 (IL-8) release in T84 cells (33).

During infection, *E. coli* is subject to different environmental conditions, for example, temperature changes that occur both in external ambient and within host. In our proteomic analysis, DnaK, GroEL and GroES were detected among the most abundant proteins (Table 1). Different studies have shown that these proteins contribute to the resistance process of EHEC under elevated temperature (34, 35). In addition, Kudva et al. (36) demonstrated that DnaK and GroEL were induced when EHEC was grown in bovine rumen fluid, thus showing the contribution of these proteins in the adaptation of EHEC to the bovine rumen.

Other type of stress commonly found by EHEC during the infection process is oxidative stress, which is generated by reactive oxygen species (ROS) such as superoxide anion (O_2_^-^) hydrogen peroxide (H_2_O_2_) and the hydroxyl radical (OH^⋅^) produced mainly by host immune response (37). Thus, to adapt and survive under this stress condition, this bacterium presents different anti-oxidant systems. We detected two members of peroxiredoxins (Prxs) family: periplasmic thiol peroxidase (Tpx) and alkyl hydroperoxide reductase C (AhpC) system (Table 1). These two antioxidant systems play an important role in the scavengers of H_2_O_2_ and organic hydroperoxides (38, 39). Glutaredoxin 3 (Grx3) was also among the most abundant proteins (Table 1). Grx3 is associated with Glutaredoxin (Grx) system, whose function is to reduce disulfide bond in target proteins to control the intracellular redox environment (40). In addition, Smirnova et al. (41) showed that glutaredoxin proteins might be involved in the resistance of E. coli to antibiotics as ampicillin. Altogether, these different systems promote an efficient pathway of antioxidant defense in EHEC that contributes to the pathogenesis of this bacterium.

The *ter* operon related to tellurite resistance is widely spread in several Gram positive and Gram negative pathogenic species (42, 43 In EDL933, this operon is composed by six genes (terZABCDE). Among the proteins expressed by that operon, only TerC was absent from our proteomic analysis. Interestingly, TerD and TerE proteins were among the most abundant proteins of EHEC O157:H7 proteome (Table 1). A study performed with Uropathogenic *E. coli* (UPEC) isolates showed that the introduction of the *ter* gene cluster contributes to improve bacterial fitness inside macrophages (44). On the other hand, Yin et al. (45) demonstrated that *ter* genes contribute to adherence of EHEC O157:H7 to epithelial cells. However, the true role of these genes in the EHEC pathogenesis remains unclear. Although tellurium is absent from the EHEC niche, interestingly, proteomic studies have detected tellurium resistance proteins in EHEC O157 proteome in different media and growth conditions such as D-MEM (46), minimal medium (47), CHROMagar™STEC (48), bovine fluid rumen (36) and under conditions that stimulate the quorum sense pathway (49). Despite the several studies in this area, more efforts are necessary to unveil the true role of the tellurium resistance proteins in EHEC pathogenesis.

### Locus of Enterocyte Effacement (LEE)

The LEE is a pathogenicity island of 35.6 kb that is organized into five polycistronic operons (*LEE1* to *LEE5*) and is an additional bicistronic operon of glr regulatory proteins (50, 51). LEE is related to intimate adherence of EHEC to cell host and is required for attaching and effacing (A/E) lesions, followed by the translocation of effector proteins that contribute mainly to host modulation of the immune system (52). In addition, LEE contains the genes that encode the Type III secretion system (T3SS) as well as some effectors molecules that are exported by this system. The T3SS is responsible for the translocation of effectors from within the host cell, whose are directly involved in the EHEC pathogenesis, mainly in the host modulation of the immune system (52). In this study the EHEC strains were grown in D-MEM, a medium known to induce expression of genes encoding T3SS (54). We identified 19 LEE-encoded proteins (Figure 1C, Supplemental material, Table S.2). Among these proteins, the most abundant were EspA (filamentous structure of the T3SS), EspB (pore formation and effector activity), Tir (translocated intimin receptor) and Intimin (outer membrane adhesin) (Figure 1C).

Interestingly, these proteins play an important role in the *E. coli* O157 adhesion (54, 55). On the other hand, EspA, EspB, Tir and Intimin are potential vaccine candidates against EHEC infection (56, 57). EspA, which was detected as the most abundant protein of LEE, forms a channel that connect the bacterial cytoplasm with the host cell; this exportation conduct allows the translocation of effectors from within the host cell (58). EspB together with EspD are responsible for the formation of the translocation pore and for the effector translocation of Tir (59). In addition, EspB can inhibit the interaction between myosin and actin, which promotes loss of microvilli and consequently contributes to the induction of diarrhea (60). The interaction between Tir and Intimin contributes directly to EHEC O157:H7 persistence during the infection process (61, 62). Furthermore, Tir and Intimin are involved in the modulation of host immunity. Tir might inhibit tumor necrosis factor receptor-associated factor 6 (TRAF6)-mediated by NF-κB activation (63). Instead, intimin can induce a T-helper cell type 1 response as well as to stimulate the proliferation of spleen CD4+ T lymphocytes and cells from lymphoid tissues (64, 65).

### Conclusion

In this work, we applied the quantitative proteomic (TMT)-based and emPAI analyses to estimate the quantification of EHEC O157:H7 proteome of combined proteomes of two EHEC O157:H7 isolates from Argentinian cattle and of the standard strainEDL933. These comprehensive proteomic analyses generated a quantitative dataset of EHEC proteome composed of a subset of proteins involved in different biological processes. All these proteins together might form a network of factors that play an important role in the pathogenesis and physiology of this pathogen. Altogether, the results presented in this study provide insights into the functional genome of EHEC O157:H7 at the protein level and could contribute to the understating of the factors associated with the biology of this pathogen.

## MATERIAL AND METHODS

### Bacterial strain and growth conditions

The EHEC O157:H7 strains Rafaela II (clade 8) and 7.1 Anguil (clade 6) isolated from cattle in Argentina and EDL933 (clade 3) strain recovered from a patient in USA were routinely maintained in Luria-Bertani broth (LB, Difco Laboratories, USA) or in LB 1.5% bacteriological agar plates, at 37°C. For the proteomic studies, bacterial strains were cultured as previously described by Amigo et al. (7). Overnight cultures of the different EHEC O157:H7 strains growth in LB were inoculated (1:50) in Dulbecco’s modified Eagle’s medium (DMEM)-F12 nutrient until reach the mid-exponential growth phase (OD_600nm_=0.6) under a 5% CO_2_ atmosphere at 37°C.

### Protein extraction and preparation of whole bacterial lysates for LC-MS/MS

After bacterial growth, protein extractions were performed according to Amigo et al. (7). Three biological replicates of each culture were centrifuged at 5000 × g for 20 min at 4°C. The cell pellets were resuspended in ice-cold lysis buffer (50 mM Tris-HCl, pH 7.5, 25 mM NaCl, 5 mM DTT and 1 mM PMSF) and disrupted by three cycles in liquid N2 and subsequently placed in boiling water. The resulting lysates were centrifuged at 30,000 × g for 10 min and precipitated with 5 volumes of ice-cold acetone at −20°C overnight. Next, the protein pellets were resuspended in buffer containing 8 M urea, 2 M thiocarbamide and 200 mM tetraethylammonium bromide at pH 8.5. The protein concentration was determined by the Bradford assay using BSA curve as a standard. Subsequently, the samples were reduced with tris-(2-carboxyethyl)-phosphine (200 mM), alkylated with iodoacetamide (375 mM) and enzymatically digested with sequencing grade trypsin. Finally, the samples were labeled with TMT Reagents 6-plex Kit according to the manufacturer’s instructions.

### Liquid chromatography and mass spectrometry

The proteomic analyses were performed using High pH Reverse Phase Fractionation and Nano LC-MS/MS Analysis by Orbitrap Fusion. Firstly, the labeled peptides were pooled together and desalted using Sep-Pak SPE (Waters) to remove salt ions. The hpRP chromatography was performed with Dionex UltiMate 3000 model on an Xterra MS C18 column (3.5 um, 2.1 × 150 mm, Waters). The sample were dissolved in buffer A (20 mM ammonium formate, pH 9.5) and then eluted with a gradient of 10 to 45% buffer B (80% acetonitrile (ACN)/20% 20 mM NH_4_HCO_2_) for 30 min, followed by 45% to 90% buffer B for 10 min, and a 5-min hold at 90% buffer B. Forty-eight fractions collected at 1 min intervals were merged into 12 fractions. The nano LC MS/MS analysis was carried out using a Orbitrap Fusion tribrid (Thermo-Fisher Scientific, San Jose, CA) mass spectrometer with an UltiMate 3000 RSLC nano system (Thermo-Dionex, Sunnyvale, CA). The fraction was injected onto a PepMap C18 trapping column (5 μm, 200 μm × 1 cm, Dionex) and separated on a PepMap C18 RP nano column (3 μm, 75 μm × 15 cm, Dionex). For all the analysis, the mass spectrometer was operated in positive ion mode, MS spectra were acquired across 350–1550 m/z scan mass range, at a resolution of 12,0000 in the Orbitrap with the max injection time of 50 ms. Tandem mass spectra were recorded in high sensitivity mode (resolution >30000) and made by HCD at normalized collision energy of 40. Each cycle of data-dependent acquisition (DDA) mode selected the top10 most intense peaks for fragmentation. The data were acquired with Xcalibur 2.1 software (Thermo-Fisher Scientific).

### Database searching, Protein Identification and Abundance Estimation

Analyses were carried out by Mascot (version 2.4.1, Matrix Science, Boston, MA) against the databases described below. The raw files from MS/MS datasets were searched against a combined database of proteins composed by the annotation of EHEC O157:H7 TW14359 and EHEC O157:H7 EDL933 genomes. For protein identification, the parameters were used as follows: one missed cleavage was allowed with fixed carbamidomethylation (Cys), fixed 6-plex TMT modifications on Lys and N-terminal amines and variable modifications of oxidation (Met), deamidation (Asn and Gln). The peptide and fragment mass tolerance values were set as 8 ppm and 20 millimass units (mmu), respectively. The target-decoy strategy (66) and the Mascot-integrated percolator calculation were applied to estimate the false discovery rate (FDR). Only peptides above "identity" were counted as identified. Furthermore, a protein must produce at least two unique peptides that generate a complete TMT reporter ion series to be confidently quantified. MS/MS based peptide and protein identifications were validated via Scaffold (version Scaffold_4.4.3, Proteome Software Inc., Portland, OR).

Peptide identifications were accepted when the peptide FDR is below 1.0%. Protein identifications were accepted when the protein FDR is below 1.0% and at least two unique peptides could be quantified. The proteins that contained similar peptides and that could not be differentiated based on MS/MS analysis alone were grouped to satisfy the principles of parsimony. The intensities of reporter ions for each valid spectrum were normalized. The reference channels were normalized to produce a 1:1-fold change. All normalization calculations were performed using medians. The protein abundance index was obtained by emPAI analysis using protein identification data from Mascot and Scaffold and used to calculate the emPAI algorithm. The equation emPAI/Σ (emPAI) × 100 was used to calculate the protein content in mol%.

### Bioinformatics analysis

Functional annotations were assigned by the COG database (14). Metabolic pathways were determined by analyzing proteins with the Kyoto Encyclopedia of Genes pathways and Genomes (KEGG) (19).

## SUPPLEMENTAL MATERIAL

Supplemental File

## ACKNOWLEDGMENTS

The present study was supported by grants PICT #0211 from FONCYT, Argentina, PNBIO1131034 INTA, Argentina; Presidential Foundation of Guangdong Academy of Agricultural Sciences (No.: 201320), and Science and Technology Program of Guangdong Province, (2016B070701013) China.

WMdS, AA, PV, ML and AC are CONICET fellows. NA holds a CONICET fellowship. We thank Julia Sabio y García for her critical reading.

